# Real Time and Delayed Effects of Subcortical Low Intensity Focused Ultrasound

**DOI:** 10.1101/2020.09.05.283747

**Authors:** Joshua A. Cain, Shakthi Visagan, Micah A. Johnson, Julia Crone, Robin Blades, Norman M. Spivak, David W. Shattuck, Martin M. Monti

## Abstract

Deep brain nuclei are integral components of large-scale circuits mediating important cognitive and sensorimotor functions. However, because they fall outside the domain of conventional non-invasive neuromodulatory techniques, their study has been primarily based on neuropsychological models, limiting the ability to fully characterize their role and to develop interventions in cases where they are damaged. To address this gap, we used the emerging technology of non-invasive low-intensity focused ultrasound (LIFU) to directly modulate left lateralized basal ganglia structures in healthy volunteers. During sonication, we observed local and distal decreases in blood oxygenation level dependent (BOLD) signal in the targeted left globus pallidus (GP) and in large-scale cortical networks. We also observed a generalized decrease in relative perfusion throughout the cerebrum following sonication. These results show, for the first time using functional MRI data, the ability to modulate deep-brain nuclei using LIFU while measuring its local and global consequences, opening the door for future applications of subcortical LIFU.

## Introduction

While routinely used to modulate the cortex non-invasively, established neuromodulatory techniques such as transcranial magnetic stimulation (TMS) and transcranial electrical stimulation (tES) remain limited in both the domains of spatial precision and the depth of their influence^1,2^. Because such non-invasive protocols are unable to selectively target structures below the cortical mantle, reversible neuromodulation of the deep brain has been limited to invasive techniques, such as deep brain stimulation (DBS), which involve the surgical implantation of electrodes and are thus limited to (severe) patient populations. However, the emerging technology of low-intensity focused ultrasound (LIFU) has increasingly been shown to address this gap; indeed, LIFU has demonstrated the capacity to both inhibit as well as excite subcortical tissues safely and reversibly in model organisms (i.e., rats ^3–7^, pigs ^8^, and macaques^9^, as well as, recently, healthy human subjects^10^) with a spatial precision far exceeding that of TMS or tES^1,2,8,11,12^.

The ability of LIFU to selectively modulate subcortical tissue non-invasively potentially makes way for causal inferences in the study of subcortical networks as well as the treatment of many neurological conditions. The lentiform nuclei are of particular interest for their role in a cortico-basal ganglia-cortical circuit ostensibly mediating motor refinement, cognitive functioning, and arousal^13,14^. Moreover, these structures are relatively accessible to LIFU through the temporal window, the thinnest section of the temporal bone and, thus, the ideal cranial entry-point for minimizing ultrasound attenuation and refraction through skull when utilizing a single-element transducer. Despite the centrality of basal ganglia-cortical circuits to several aspects of human cognition^13^, knowledge of their precise structure and function remains incomplete, yet evolving. Only recently, for instance, has the field begun appreciating the contribution of the lentiform nuclei to maintaining electro-cortical and/or behavioral arousal^15,16^, as evidenced by both animal models and human studies, putatively through a recently identified extra-thalamic direct pallido-cortical pathway^17–19^. LIFU thus promises the unprecedented ability of performing causal investigation into the role of these circuits in healthy volunteers. While clinical translation of this technique is already ongoing^20^ in the context of Disorders of Consciousness^14^(DOC) after severe brain injury, pallidal LIFU is likely to be applicable to other conditions such as obsessive-compulsive disorder^21^, Tourette syndrome^22^, treatment-resistant depression,^23^ Huntington’s disease, and Parkinson’s disease.^24,25^

Here, we administered two sessions of deep-brain LIFU in 16 healthy volunteers. In each session we delivered two 5-minute doses of LIFU aimed at the left globus pallidus. Each dose was administered in 30-second blocks separated by 30-second rest intervals. Given the current uncertainty with respect to how different sonication parameters (e.g., duty cycle, pulse repetition frequency, tone burst duration) relate to local and distal brain modulation, two sonication modes modeled after prior work^4–7^ were administered, one per session. Online (during sonication) brain responses to each sonication were assessed with T2*-weighted blood oxygenation level dependent (BOLD) signal. Offline (post-sonication) effects were assessed before and after sonication with perfusion-weighted arterial spin labeling (ASL). The main aims of the work were to assess the online and offline topography (local and global) and valence (i.e., up-/down-modulation) of the brain response to each sonication setting.

## Results

We present the results of this work in three main sections. First, we describe our sonication settings as well as the spatial characteristics of our ultrasound beam when passed through free water (measured empirically) and bone (simulated using k-wave^26^ in Matlab). Second, we report online local and global effects of LIFU sonication with an ROI analysis of the principal target (left GP) and proximal structures (i.e., left putamen and left thalamus), as well as a full brain analysis of the same BOLD data. Finally, we describe offline effects of LIFU both locally, with an ROI analysis of ASL perfusion data, and globally, with a full-brain analysis of the same perfusion data.

### Ultrasound Waveform Through Water and Bone

Previous simulations of ultrasound propagation have demonstrated the general maintenance of focal shape and focal location when passing through the human skull, including through the temporal bone^28^. To ensure that these results generalize to our LIFU parameters and our trajectory, we have simulated, in three dimensions, 5ms (equivalent to 1 pulse in LIFU Mode 2) of LIFU propagation at our parameters through the temporal bone of a high-resolution CT (visible human^®29^) and into the left GP. Regarding refraction, the point of maximum pressure of this simulation, compared to simulation through water alone, deviated by 0.93 cm, in the range of previous findings^28,30^. As depicted in Figure 2 C and D, our results suggest that energy deposition remains inside the targeted left GP while not significantly impacting ventral or dorsal structures (e.g., left hippocampus), despite some expected deviation. The thresholded (−50%) pattern of energy deposition subsumes portions of the left thalamus and left putamen, supporting the attention given here to these structures (Figure 2 D, Cyan). Regarding attenuation, skull bone attenuated the energy reaching the brain significantly, as expected, resulting in a peak in-brain pressure 12.35% of that simulated in water alone. It is important to note that the depictions of our ultrasound beam here, for both water and bone simulations, are thresholded at an arbitrary value: (−50%) from maximal in-brain pressure. While the skull deforms and “flattens” the energy deposition to some extent, resulting in a larger area exceeding that −50% threshold (see Figure 2 B), an oblong focus of high intensity remains with a degree of refraction (0.93cm) that retains energy deposition into pallidal tissues. Note that cumulative energy deposition over time was recorded at the point of max intensity and is highly linear (see Figure S8), suggesting that these results also generalize to the shorter 0.5ms pulse.

## Region of Interest (ROI) Analysis

### BOLD – ROI

We assessed the local influence of sonication during sonication using both parameter sets on BOLD signal as compared to baseline (30 second inter-sonication blocks when no ultrasound is applied). A 2 x 2 x 3 repeated measures ANOVA (as well as its Bayesian equivalent) was performed with Parameter Set (Mode 1 or Mode 2), Run (Sonication 1 or Sonication 2 within session), and ROI (Left Putamen, Left GP, Left Thalamus) as factors; this revealed a significant effect of Parameter, *F*(1,15) = 5.442, *p* = .034, BF_Inclusion_ = 78.620. A significant interaction (in the frequentist but not Bayesian approach) between Parameter Set, Run, and ROI was found *F*(1,15) = 9.055, *p* = 8.36e^-4^, BF_Inclusion_ = 0.6 47). Based on this finding as well as our a-priori expectation of regional effects from LIFU (see Discussion), a follow-up two-way ANOVA was performed for each ROI. While no significant effects were found for the Left Putamen, a significant effect of parameter was found in both the Left GP (*F*(1,15) = 4.585, *p* = .049, BFinclusion = 2.49) as well as the Left Thalamus *F*(1,15) = 5.115, *p* = .039, BFinclusion = 1.53) with reduced BOLD found during sonication in LIFU Mode 1 as compared to LIFU Mode 2 for both ROIs. To assess if either parameter set induced a change in BOLD signal from baseline, marginal means were assessed for each parameter set for each ROI. A reduction in BOLD from baseline during sonication in Mode 1 was found in the LEFT GP, *t*(15) = −2.923, *p_Šidák_* = .013, BF_10_ = 5.54, and the Left Thalamus, *t*(15) = −2.436, *p_Šidák_* = .042, BF_10_ = 3.62. These results suggest that sonication in Mode 1 significantly reduced BOLD signal in the Left GP and the adjacent Left Thalamus when compared to sonication in Mode 2 and when compared to baseline. Sonication in Mode 2, despite its identical intensity and DC of 5% as compared to Mode 1, induced no significant influences on BOLD in these regions.

### ASL – ROI

To assess longer-term effects of pallidal LIFU on relative brain perfusion, we performed a three-way repeated measured ANOVA, as well as its Bayesian equivalent, including sonication parameter (Mode 1, Mode 2), time point (Pre LIFU 1, Post LIFU 1, Post LIFU 2) and ROI (Left Putamen, Left GP, Left Thalamus) as factors. As depicted in Figure 3, we only observed a main effect of time point (*F_Greenhouse-Geisser_* (1.303, 19.547) = 7.926, *p* = .007; BF_Inclusion_ = 10.174). Post-hoc analysis revealed that this effect was driven by a large decrease in perfusion, within the ROIs, following LIFU 1 (*t*(15) = 3.231, *p_holm_* = .006; BF_10_ = 138.915), with no additional decrease observed following LIFU 2 (*t*(15) = .399, *p* = .693; BF_10_ = 0.175). No main effect of sonication parameter was found (*F_Greenhouse-Geisser_*(1,15) = .177, *p* = .680; BF_Inclusion_ = 0.164). An interaction between time-point and ROI was also observed (*F_Greenhouse-Geisser_* (2.290, 34.353) = 3.180, *p* = .048) albeit with weak support (BF_Inclusion_ = 0.055) consistent with the fact that follow-up 2-way repeated measures ANOVAs, one per ROI, generally revealed the same pattern of decreased relative perfusion over time seen in the 3-way analysis.

In the Left Putamen, a main effect of Run was found (*F_Greenhouse-Geisser_* (1.396, 20.102) = 4.873, *p* = .028; BF_Inclusion_ = 0.824). Follow up revealed a more linear trend with no significant difference between run 1 and 2 *t*(15) = 2.537, *p_holm_* = .074; BF_10_ = 0.654) or between run 2 and 3, (*t*(15) = .840, *p_holm_* = .407; BF_10_ = 1.325). In the Left GP, a main effect of Run was found (*F_Greenhouse-Geisser_* (1.141, 18.739) = 7.543, *p* = .011; BF_Inclusion_ = 1.719). Follow up revealed a significant difference between run 1 and 2 was found 2 (*t*(15) = 2.867, *p_holm_* = .015; BF_10_ = 1.159) with no difference between run 2 and 3 found. In the Left Thalamus, a main effect of Run was found (*F_Greenhouse-Geisser_* (1.629, 21.512) = 7.816, *p* = .004; BF_Inclusion_ = 3.707). Follow up revealed a significant difference between run 1 and 2 (*t*(15) = 3.561, *p_holm_* = .004; BF_10_ = 8.242) with no difference between run 2 and 3 found.

## Whole Brain Analysis

### BOLD – Whole Brain

Full-brain analysis of the BOLD response during LIFU sonication revealed several foci of reduced BOLD response during Mode 1 (PRF = 100Hz, PW = 0.5ms; see Figure 4). Significant clusters included right and left pre- and post-central gyri, frontal polar cortex, posterior cingulate cortex, and Heschl’s Gyrus (see also Table S1). Given the known conservative bias of FSL-FLAME 1+2^31,32^ in single-sample t-tests at the employed cluster defining threshold (CDT; equivalent to p=0.001; see Figure 1A,B in ref 31^31^), results are shown at two CDT values (Figure 4; p=0.001, in violet, and p=0.005, in blue). At this lower threshold, an additional cluster is visible in the medial frontal cortex and significant clusters expand to subsume portions of the dorsal thalamus. No area of increased BOLD response was observed at either CDT during LIFU Mode 1 sonication. Furthermore, no significant foci of increased or decreased BOLD signal were observed during LIFU Mode 2 (PRF = 10Hz, PW = 5ms) sonication at either CDT.

**Figure 1.**
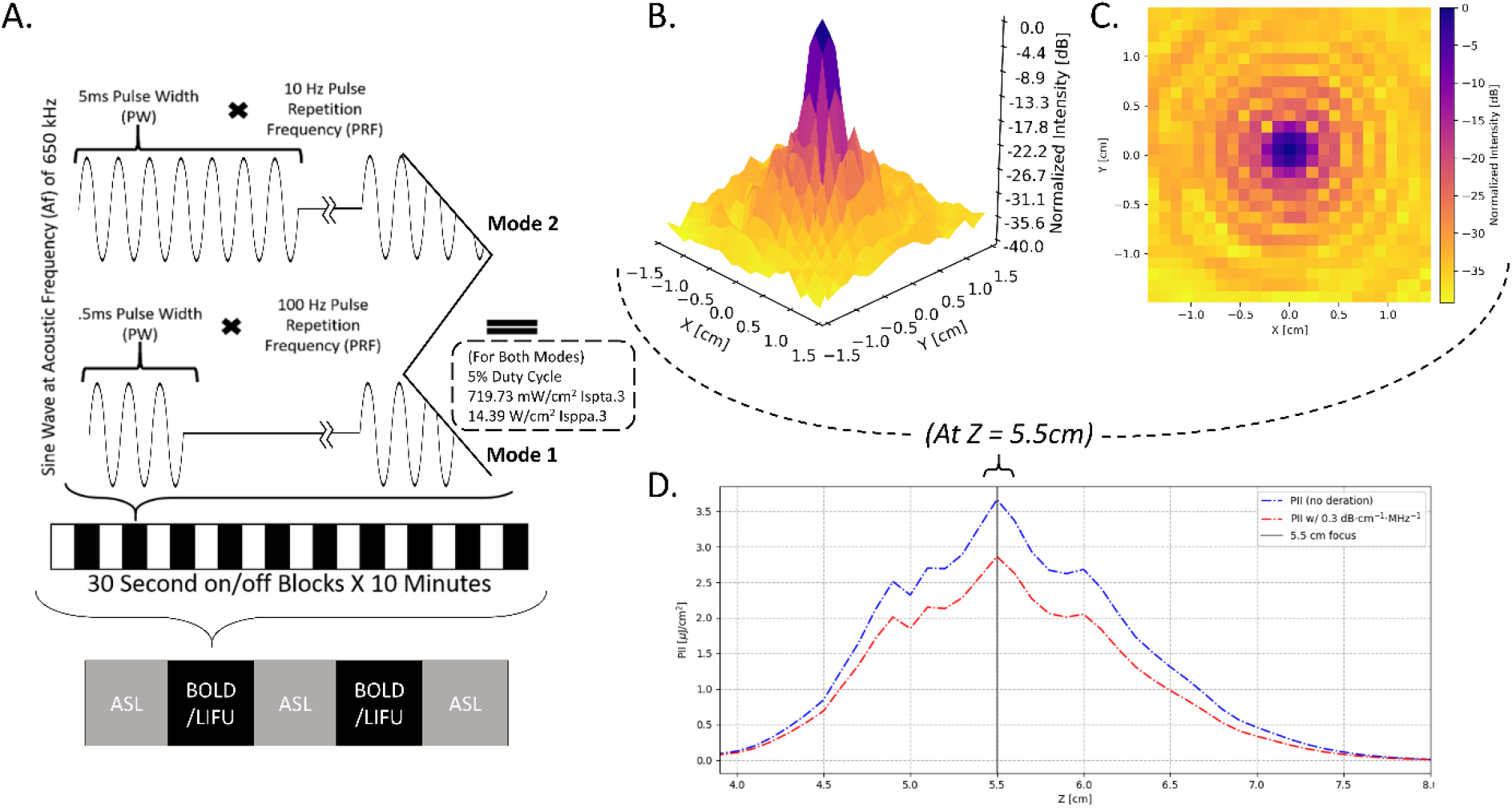
Sonication Parameters, Experimental Design, and Water Tank Measurements of Beam Properties. **A)** Parameters for each LIFU mode utilized including pulsing schedule, block design, and intensity. I_sppa.3_ = Spatial Peak Pulse Average Intensity. I_spta.3_ = Spatial Peak Temporal Average Intensity; “.3” denotes deration (attenuated intensity at 0.3 dB/cm-MHz) through human tissue. Here, we have applied LIFU in two sessions, utilizing different parameter sets in each with LIFU Mode 1 having a pulse repetition frequency (PRF) of 100Hz PRF and a pulse width (PW) of 0.5ms PW while LIFU Mode 2 = 10Hz PRF, 5ms PW; all other factors including duty cycle (DC) = 5% and intensity I_spta.3_ (Spatial Peak Temporal Average)^27^ = 720 mW/cm^2^; I_sppa.3_ (Spatial Peak Pulse Average)^27^ = 14.40 W/cm^2^ were held constant. **B,C)** Intensity in the radial plane (X/Y plane, extending from focal point of ultrasound beam 5.5cm from transducer surface) shown in both 3 (**B**) and 2 (**C**) dimensions. A 50% (−3dB) reduction in peak intensity occurs in an area approximately 0.5 cm in width. Note that the decibel scale is nonlinear and −3 dB approximately corresponds to a 50% reduction in intensity; this scale is normalized to maximal intensity, where peak intensity equals 0dB. **D)** Intensity in the longitudinal plane (Z plane, extending from transducer) in absolute (pulse intensity integral (PII); “.3”denoting absorption in human tissue at 0.3 dB/cm-MHz) values of Z correspond to distance from the transducer surface. Note the peak intensity 5.5 cm from the transducer surface and that a 50% (−3dB) reduction in peak intensity is found in an area approximately 1.5 cm in length.

**Figure 2:**
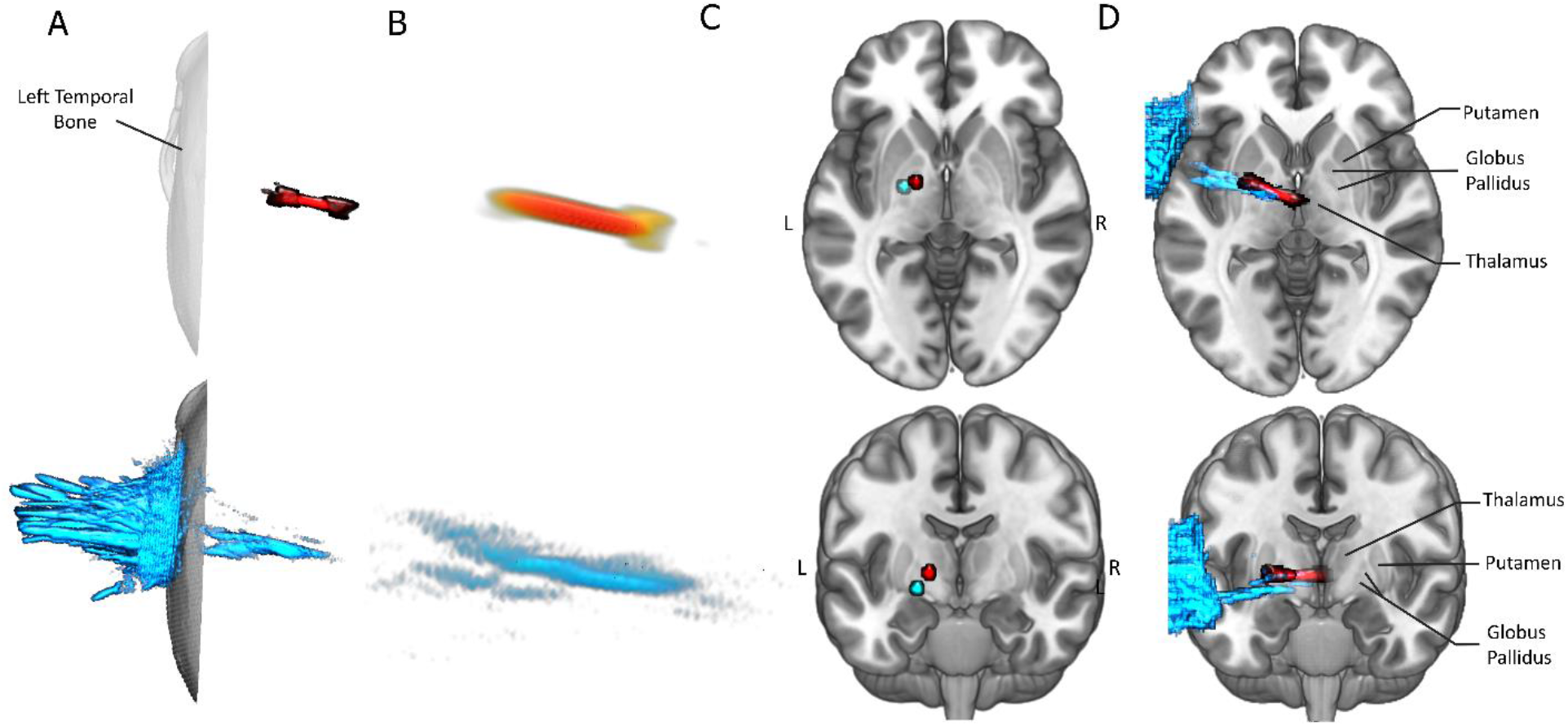
Numerical Modeling through Bone and Water. Here, one 5ms pulse (corresponding to 1 pulse of Mode 2) of our ultrasound beam is simulated twice, in water (red), and through the temporal bone of a human computed tomography (CT) image (cyan). The same trajectory, which targets the left GP, was used in both simulations. The maximal pressure for each voxel over the course of the simulation is visualized. Only Voxels exceeding 50% (−3dB) of the maximum in-brain pressure are presented. **A)** Depiction of the effect of bone on beam shape and position. Note that bone (bottom, cyan) appears to flatten, deform, and laterally retract the ultrasound beam compared to the water condition (top, red). However, the general expression of an elongated beam and its general location is retained. Note that most energy (A bottom) is deposited into and reflected off bone when it is present. **B)** Higher-resolution depiction of ultrasound beam in water (B top) and through bone (B bottom). **C)** Location of maximum pressures following propagation through water (red) and through bone (cyan). Points were mapped into MNI space for visualization. Note that both reside inside the left pallidal target. While the effect of bone moves the peak pressure somewhat ventral and lateral, the total translation is 0.93 cm. **D)** Depiction of whole ultrasound beam through water (cyan) and bone (red). Simulations were mapped into MNI space for visualization. Note that energy deposition into portions of the left lentiform nuclei and left thalamus exceeded the threshold of −50% maximum pressure.

**Figure 3:**
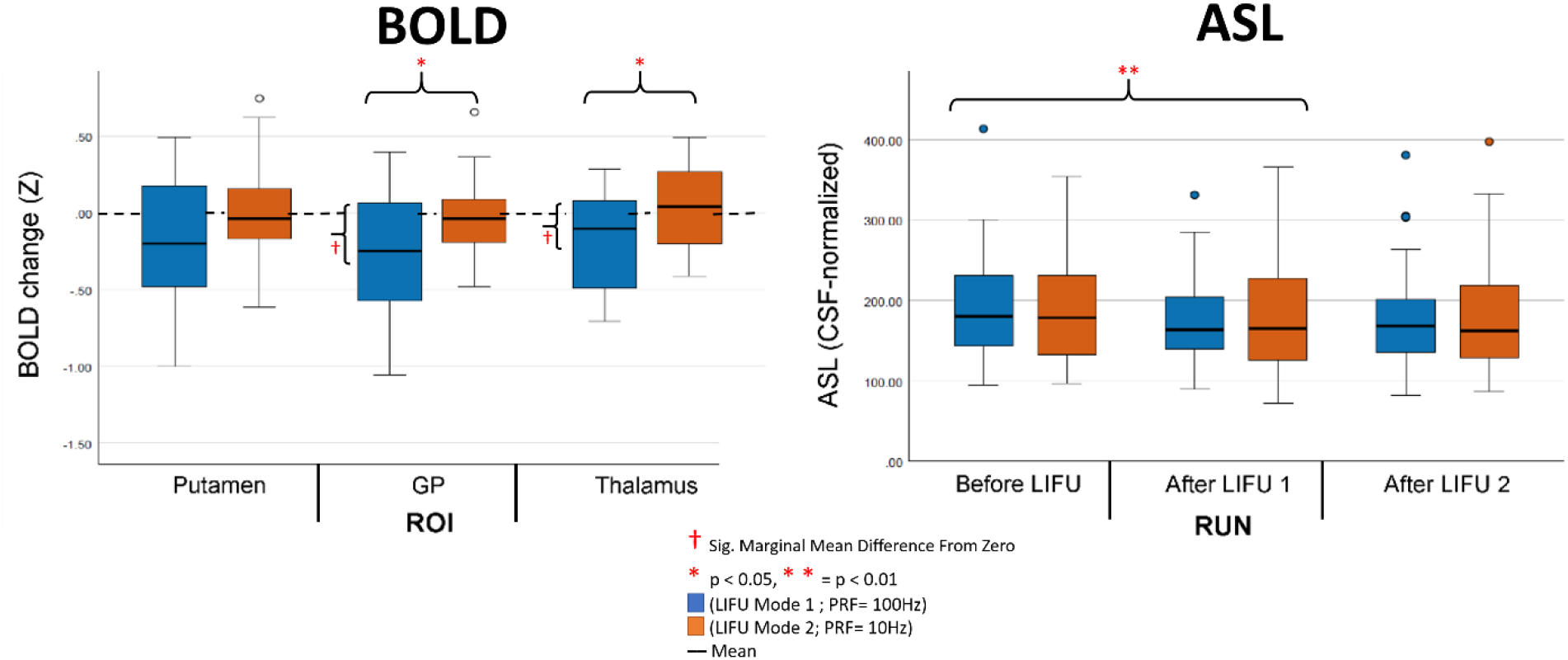
ROI analysis results. Left: online changes in BOLD signal during LIFU sonication blocks compared to inter-sonication blocks (i.e., baseline) for the target Left GP ROI and in the proximal Left putamen and Left thalamus ROIs during Mode 1 (blue) and Mode 2 (orange) sonication, compared to baseline (Red crosses indicate a significant difference from baseline for an individual condition while red Asterisks indicate significant difference across conditions). Right: offline changes in ROI perfusion before LIFU, after sonication 1, and after sonication 2 for each sonication mode (Red Asterisks indicate significant difference across conditions). Whiskers represent 1.5 times the interquartile range above the 75^th^ percentile and below the 25^th^ percentile.

**Figure 4:**
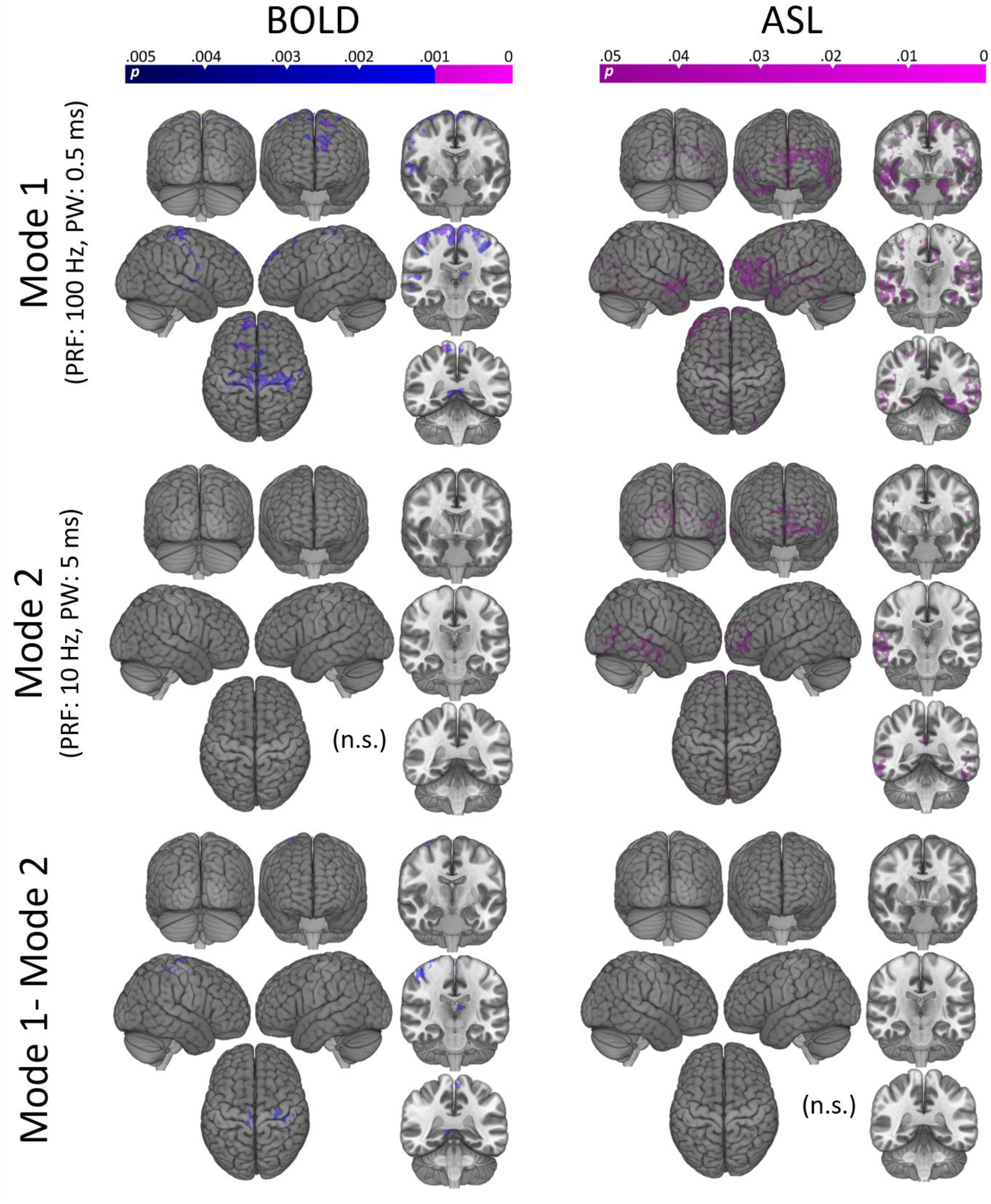
Whole brain results. Left: Regions of significant decrease in BOLD signal during sonication Mode 1 (top) and Mode 2 (middle) compared to inter-sonication periods (i.e., baseline). Subtraction of these results are also shown (bottom). Statistical maps were obtained using a mixed effects model (FLAME 1+2) as implemented in FSL^34^, and are shown at two levels of cluster correction for multiplicity (CDT set at p < 0.005, in blue, and at p < 0.001 in violet). Significant reduction in BOLD is found in several cortical regions, at both thresholds, for Mode 1. No regions of increased BOLD signal were observed for Mode 1. No regional increase or decrease was observed for Mode 2. Right: Regions of significant decrease in relative perfusion after sonication in Mode 1 (top) and Mode 2 (middle). Subtraction of these results are also shown (bottom). Statistical maps were obtained with a non-parametric approach (FSL randomise), as implemented in FSL, and are here shown at a level of p <0.05 corrected for multiplicity with threshold free cluster enhancement (TFCE). No significant increase in ASL was observed for either sonication mode.

Direct comparison of the BOLD response under the two stimulation modes revealed no significant differences at a CDT of p <.001. When displayed at the lower CDT (Figure 4 bottom left), however, some foci of significant difference were observed in the very foci reported for Mode 1 (pre- and post-central gyri and posterior cingulate cortex/ left dorsal thalamus).

### ASL – Whole Brain

Given the ROI result, full-brain relative perfusion images were analyzed by contrasting the average brain perfusion after LIFU (i.e., the average of time-point 2 and time-point 3) to the relative perfusion prior to LIFU (as shown in figure S4, the results are qualitatively unchanged when the data were analyzed with a linear model [i.e., time-point 3 versus time-point 1]). As shown in Figure 4, a nonparametric permutation t-test, corrected for multiple comparisons with threshold-free cluster enhancement, TFCE,^33^ revealed broad decrease in perfusion throughout the cerebrum for both LIFU modes. In accordance with our ROI results, no significant difference was found when directly comparing sonication modes, and no increase in perfusion was found for either mode.

## Discussion

This study is the first demonstration of a group-wide^35^ MRI response from subcortical focused ultrasound in humans. Taken together, our results suggest that, when targeting the GP and adjacent structures: **1)** LIFU, across two modalities (i.e., BOLD, ASL), in Mode 1 appears neuroactive both during (i.e., online) and after (i.e., offline) sonication. **2)** The valence of online, offline, local, and distal effects all appear inhibitory. **3)** LIFU parameters (PRF and PW) indeed appear to impact the acute effect of LIFU (on the order of 30s, as observed with BOLD), while statistically similar changes in perfusion suggest long-term effects (on the order of minutes, as observed with ASL) may not be as susceptible to these differences, perhaps adding a temporal dimension to the parameter-space of LIFU.

Firstly, we asked if online LIFU could modulate BOLD signal in the target of interest as well as throughout the brain. We found an apparent inhibition of our intended target nuclei, as well as the adjacent left thalamus, during 30-second trains of LIFU in Mode 1 but not in Mode 2 (see Figure 3). At the whole-brain level we also found an online inhibitory effect of LIFU in Mode 1, but not Mode 2, within the primary somatomotor cortex, left dorsal thalamus, as well as frontal and posterior cingulate association cortex concurrent with pallidal sonication (see Figure 4, Table S1). It remains to be seen if LIFU induces changes in neurovascular coupling irrespective of changes in neural activity; certainly, higher intensity ultrasound may cause vasoconstriction^36^. This could theoretically explain BOLD changes in our targeted structures but is much less relevant to our findings distal to the targeted nuclei. Regardless, the absence of any BOLD response in Mode 2 either locally or distally provides a negative control, suggesting that our results are specific to the parameters employed and argues against the notion that these results stem from artifact (i.e., neurovascular, MRI^35^, and/or auditory^37^) or from inflated type 1 error^31^.

While GP connectivity remains elusive, the results presented here demonstrate a plausible pattern of effects. Inhibition of the left pallidum significantly affects all other components of the cortico-basal ganglia-thalamo-cortical circuit underlying motor refinement^13^ and cognitive functioning^14^, either by way of thalamus as conduit^14^ or through direct pallido-cortical projections^17–19,38^. Our BOLD results—primarily found in frontoparietal cortex and thalamic tissues—fit within this framework. Recording activity from both the target of interest and globally is key to understanding the valence of LIFU’s direct influence on neural tissue and the network effects this may bring about. Despite this, direct measurement from targeted subcortical structures is rare in LIFU studies (a notable exception being a recent study in macaques^9^) while it is completely absent in humans until this point, rather being inferred from downstream impacts on cortical activity^5,8,10^. Dual local and distal effects found here bolster the notion that fMRI can be used to effectively understand LIFU and its parameter space in healthy human subjects. On the contrary, only distal effects could have been found, which is often the case in TMS-fMRI^39^, and which would have severely impaired valence-specific conclusions about LIFU’s direct influence. These results, which are based on a the contrast of 30s blocks of LIFU compared to 30s inter-sonication baseline blocks, suggest that the effects of LIFU can vary over a time-course of seconds, despite our own findings in relative perfusion as well as prior results demonstrating sustained influence from LIFU lasting minutes^7^ or hours^9,40,41^. This finding supports the feasibility of classical block designs in future studies and invites investigation of online behavioral impacts from LIFU. Furthermore, our findings of pallidal as well as cortical (most relevantly in the primary somatomotor cortex) effects of pallidal LIFU also mirror the results of studies using contemporaneous neuroimaging and pallidal DBS^42,43^, potentially suggesting that this non-invasive technique might be employed in the future to evaluate patient suitability for invasive stimulatory procedures or as a treatment intervention itself.

In addition, we used ASL to characterize the relative longer-term effects of our sonication (i.e., minutes, rather than seconds following cessation). Unlike BOLD, which is derived from more complex interactions between blood flow and acute neural activity, ASL quantifies cerebral blood perfusion alone. This reduces the inter-subject and inter-trial variability of ASL relative to BOLD^44^, making ASL preferable for detecting disparate neural activity between distant time points. Here, we again found an apparent inhibition (decreased perfusion) of our target nuclei (and the adjacent putamen, thalamus) in the minutes following sonication. Again, this inhibition was extended to the cortex, where decreased perfusion was found throughout the cerebrum (see Figure 4). However, here, we find the same pattern of results following sonication in both Mode 1 and Mode 2.

These results corroborate other findings of an offline influence from LIFU^8,9,12,40,41^. However, the estimated focal intensities used here are approximately an order of magnitude lower than recent studies in pigs^8^ and macaques^9,40,41^. The time course of LIFU’s influence on neural activity remains unclear and appears to vary dramatically between experimental paradigms, ranging from seconds^45^ to over an hour^9,40,41^ with a trend towards long-term, offline effects occurring following longer sonication periods^12^. An offline impact may be expected here given that the 10 minutes of total sonication utilized far exceeded the 40 seconds necessary to elicit effects in the tens of minutes to hour range^9,40,41^, despite the lower intensities used here. A significant perfusion effect of LIFU in Mode 2 (PRF = 10Hz) here despite null effects on BOLD suggests either the heightened discriminability of ASL compared to BOLD^44^ in this context or that modifying LIFU parameters may impact the time-course of LIFU’s influence. Given the recent offline behavioral effects found in non-human primates^41^, these results further support investigation into offline behavioral effects in humans at these intensities more comfortably within the range deemed safe for ultrasound imaging of the human cranium^27^. A growing consensus surrounding the offline impact of LIFU supports its potential use in clinical applications seeking persistent impacts on neural activity^20,46^. However, these early results should be replicated— ASL measures, like BOLD, may be confounded by changes in physiological arousal^47^ and its consequences (i.e., blood CO2 content^48^, and blood pressure^49^), as well as any as-yet undiscovered direct impacts of LIFU on neurovascular coupling.

Others have reported differential neuromodulatory effects from LIFU applied at different PRFs when intensity is held constant^45^; however, these differences are slight in comparison to alterations of other parameters, such as intensity or duty cycle^45^. LIFU parameters such as PRF are likely to interact with duty cycle, intensity, etc. in a high dimensional parameter space that is far from fully characterized, as evidenced by divergent findings between studies despite similar parameters (e.g., apparent disruption^8^ as well as excitation^7^ at PRF = 10Hz with duty cycles near 50%). A possible explanation for these apparent contradictions, it is likely that LIFU modulation may be tissue-type selective. While the mechanisms underlying neuromodulation through LIFU remains debated, principle theories concern an interaction of mechanical pressure on neural tissue and lipid bilayer and/or membrane protein dynamics which brings about depolarization by altering membrane permeability or capacitance^50,51^. Tissue-specific effects are thus likely to be driven by differential expression of membrane proteins or the variable membrane dynamics between cell types. For instance, it has been proposed that inhibition through LIFU is achieved through preferential excitation of GABAergic interneurons rather than hyperpolarization of single cells^50,52^ and that this is the mechanism underlying a general trend of inhibition during lower duty cycles. While much more work is needed to assess how LIFU interacts with the microstructure of the basal ganglia, these results support the notion of inhibition using low duty cycles, a step towards clinical applications for LIFU in our targeted tissues which absolutely requires valent-specific interventions.

The dimensions of the ultrasound beam produced with our single-element transducer ensure energy deposition into adjacent structures when targeting the GP due to its small size. This, indeed, is a limitation (as discussed below) in the use of LIFU for basic science of the GP alone. However, this may be of benefit to scientific and clinical applications aimed at broader basal-ganglia-thalamo-cortical^13,14^ circuits, whose subcortical components are ideally positioned for LIFU modulation through the temporal bone. Apparent inhibition of these circuits’ components here, if proved consistent, may make way for clinical use, with an emphasis on the modulation of basal-ganglia-thalamo-cortical^13,14^ communication that is presumed to underlie the efficacy of pallidal DBS in treating some individuals with obsessive-compulsive disorder^21^, Tourette syndrome^22^, treatment-resistant depression^23^ and movement disorders such as Huntington’s disease and Parkinson’s disease^24,25^. These results are also highly relevant to the continued study of LIFU’s influence in Disorders of Consciousness^20^ as DOC symptomatology has also been linked to basal-ganglia-thalamo-cortical commmuncation^14^ and GP atrophy specifically^53^. The properties of focused ultrasound suggest it may represent a sensible evolutionary step in obtaining subcortical neuromodulation in these applications, avoiding, for instance, the risks inherent to DBS—including hemorrhage and infection^54^—and the non-selectivity of pharmaceuticals thought to impact this system (e.g. zolpidem and amantadine^55^). These findings represent a pioneering step towards such applications as well as an early foothold for the use of MR-guided LIFU in human neuroscience more broadly.

However, it should be noted that the MR-guided LIFU techniques used here carry with them several important limitations. As it stands, the use of a single-element LIFU transducer for modulation, while highly precise in its influence when compared to TMS, tES, is not perfectly so. A single-element array produces an oblong focus, rather than a highly-focal point of influence, in stark comparison to the more sophisticated multi-unit arrays utilized in neurosurgery^56^. Indeed, both the measured (here, < 0.5 cm wide in the X,Y by 1.5 cm long in the Z plane; threshold at −50% maximum pressure, see Figure 1) and modeled (see Figure 2) dimensions of our ultrasound beam ensure energy deposition into the adjacent nuclei of the putamen and thalamus as well as the large white-matter tracks that surround them. This inherent limitation complicates the placement of these results into canonical network models; thalamic and striatal function may be influenced both indirectly through modulated pallidal output as well as, to some degree, directly from LIFU itself and vice versa. Moreover, it cannot be ruled out that some or all of the apparent effects here result from modulation of the large white matter tracts nearby and passing through our focus (e.g. the internal capsule), a notion difficult to test with the methods utilized as BOLD signal is widely considered a less effective correlate of neural activity in white matter^57^. Proposed mechanisms of LIFU, while controversial as discussed above, are not incongruent with direct modulation of axonal projections^50,51^.

Another principle limitation in human LIFU research remains the variability in propagation and attenuation introduced by transmission through skull. Corroborating other findings^10,28,58^, numerical modeling performed here through a high-resolution CT image (visible human^®29^) suggests a general preservation of the ultrasound waveform observed in free water^10,28^. However, the degree of preservation in beam shape is likely to vary between individual sessions due to variable positioning, individual skull morphology and individual skull density^28^, the latter of which cannot be readily assessed using MRI at this time. Despite an effect found in the GP as a whole, these limitations prevent assertions concerning which aspect of the GP—Globus Pallidus Pars Externa (GPe) or Pars Interna (GPi)— we preferentially targeted, which is why we chose not to parcellate the structure further. However, as modulation of the GPe and GPi may have vastly different impacts on the circuits in which both participate, future research should investigate related methods that may selectively impact these two nuclei (e.g. multi-unit ultrasound arrays). While animal studies, even in large brained (and thick-skulled) animals^8^, suggest the general accuracy of linear targeting and human LIFU studies to date^10,58–60^ have often elected to forgo per-subject modeling of attenuation and refraction through skull, a fuller, more subject-specific, investigation of possible skull-refractory effects in naturalistic experimental settings in human subjects remains desirable going forward.

## Methods

### Participants

Participants included 16 healthy individuals (15 male; age = 18–44 years (M = 25.25; SD = 7.78)). Participants were screened for eligibility, reporting no history of neurological/psychiatric disorder or medical condition that may preclude safe entry into an MR environment. Participants were instructed to have the hair around their left temple below 0.5 inches during LIFU sessions due to the potential of hair to trap air bubbles and attenuate ultrasound. This requirement resulted in a high proportion of males. Participants received $150 compensation for taking part in the experiment. Written informed consent was obtained from all subjects according to the procedures approved by the UCLA Institutional Review Board.

### Experimental Design and MR Image Parameters

All participants underwent three discrete sessions in the 3 Tesla Siemens Prisma Magnetic Resonance Imaging (MRI) scanner at the Staglin IMHRO Center for Cognitive Neuroscience at UCLA. During **session 1)** baseline structural and functional BOLD data was collected. This included a T1-weighted structural sequence (MPRAGE, TR = 2000 ms, TE = 2.52 ms, voxel size 1 mm^3^), T2-weighted structural sequence (TR = 1500ms, TE = 104 ms, voxel size 1 mm^3^), a baseline arterial spin labeling image (TR = 4.6 ms, TE = 16.18 ms, 8 sequential slices, voxel size 1 mm^3^, inversion time = 1990ms, tag-controlled pulsed ASL (PASL), bolus duration = 700ms), a baseline functional image (T2*-weighted Gradient Recall Echo sequence, TR = 700 ms, TE = 33 ms, 1000 interleaved slices, voxel size 1 mm^3^), and a white-matter nulled T1-weighted image used to better capture the fine structure of subcortical nuclei. During **sessions 2 and 3)** LIFU was applied. Firstly, a T1-weighted MPRAGE sequence (see parameters above) was captured followed by transducer placement. ASL data was collected immediately following transducer placement (see parameters above). Sonication was administered in 30-second blocks for 10-minutes while functional BOLD data (see parameters above) were collected. ASL data were again collected immediately post-sonication while another sonication was administered during the collection of BOLD data (see parameters above) immediately thereafter for a total of 2, 10-minute rounds of sonication delivered in 30 second blocks or 10-minutes of sonication total. Following Sonication 2, ASL data were again collected. Sonication parameters were always held constant within sessions but differed between Mode 1 and Mode 2 for each participant between sessions, order counterbalanced between participants.

### Ultrasound Positioning

During sessions 2 and 3, the ultrasound transducer was positioned so that its center lay on the left temple (approximately 1/3 of the distance from the corner of the left eye to the left tragus for each participant and superior 2cm). Ultrasound gel (aquasonic) was firstly applied to this region in an area subsuming the diameter of the transducer and rubbed into any hair present so that no hair permeated the gel layer in order to minimize air bubbles and ensure a smooth surface for coupling. A thin layer of gel was applied to the surface of the transducer and bubbles were similarly smoothed from this layer. The transducer was then coupled to the head with gel filling any concavity between the transducer membrane and the scalp. Two straps—one horizonal and one vertical—secured the device to the participant. Next, we acquired a rapid (95 s) T1-weighted structural sequence (TR = 1900 ms, TE = 2.2 ms, voxel size 2 mm^3^). Using a circular MR fiducial and the visible center of the transducer, reference lines were drawn using the scanner console in the transverse and coronal planes to locate the target of the LIFU beam visually in three dimensions. Adjustments to the positioning were made iteratively until the trajectory of our ultrasound beam passed through the temporal bone, through the anterior dorsal aspect of the left globus pallidus transversely as well as coronally and terminating into the left thalamus. This portion of the GP was chosen for its apparent direct structural connectivity with frontal cortex^19^ and our interest in the role of GP in cognitive functioning. The dimensions of the - 3 decibel focus of our transducer when measured in water (see Figure 1) and simulated through bone (see Figure 2), as well as the orientation of the ultrasound beam, suggests that this targeting ensures the beam consistently subsumes aspects of the left GP, with the adjacent left putamen and left thalamus also likely impacted directly by LIFU.

### Ultrasound Waveform

A BXPulsar 1001, Brainsonix Inc. ultrasound device was used in two modes. While both modes employ a 650kHz carrier wave, the distinction between them is a high PRF (100 Hz) with low Pulse Width (0.5ms) for Mode 1 and a low PRF (10Hz) with high Pulse Width (5ms) for Mode 2. Pulsation was administered in 2, 10-minute sessions on each of 2 different days and thus in 4, 10-minute sessions total. Within each 10-minute session, pulsation alternated between 30s of LIFU and 30s of no-LIFU for a total of 10, 30s trains of pulsation per sonication (5-minutes of LIFU total per 10-minute session) - a standard block design amenable to BOLD MRI. For both modes, Duty Cycle = 5%, I_spta.3_ = 720 mW/cm^2^; I_sppa.3_ = 14.40 W/cm^2^ (see Figure 1a). This intensity falls under the safety guidelines for ultrasound imaging of the cranium provided by the FDA^27^.

### Ultrasound Simulation

We simulated the propagation of the ultrasound waveform through a human skull using acoustic wave equations implemented in MATLAB using k-Wave (v1.1)^26^, a k-space pseudospectral solver toolbox. We applied these equations to computational head models derived from imaging data from the Visible Human Project^®29^, provided courtesy of the U.S. National Library of Medicine. Specifically, we used the head CT (dimensions: 512 x 512 x 512 voxels) and its paired T1-weighted MRI (dimensions: 196 x 231 x 67 voxels) data from the Visible Human^®^ Male (VHM) dataset^29^. The VHM MRI data were aligned to the VHM CT using FSL’s FLIRT program (6DOF model) and resampled to the voxel grid of the CT data. We manually defined a target beam trajectory through the temporal bone and into the left pallidum on a reference MRI brain atlas (MNI152, 1mm). This enabled improved identification of relevant anatomy for landmark selection as compared to the visible human^®^ dataset’s male MRI, which is of relatively low resolution. The trajectory was defined by selecting two anatomical points in the MNI152 atlas: 1) a target point in the anterodorsal pallidum, at which the ultrasound was aimed from the left side of the head; and 2) a point denoting the center of the transducer’s face outside the head at the point of contact with the skin. Care was taken to ensure the transducer’s membrane was flush with the skull, as was the case in our experimental setting. We registered the MNI152 atlas to the transformed VHM MRI data using FLIRT (12 DOF model), and used this transform to map the target and transducer points to the space of the VHM CT data.

Head models were derived from the CT data as follows. We first resampled the CT data into a 512×512×512 volume with isotropic voxels (0.489mm x 0.489mm x 0.489mm) using trilinear interpolation. This defined the simulation grid, whose dimensions enabled us to take advantage of the speed offered by the Fast Fourier Transform used in k-Wave’s Fourier collocation method when calculating spatial gradients. We generated three 3D arrays based on the CT scan that modeled the 1) density, 2) speed of sound, and 3) nonlinearities of the varying materials (e.g., scalp, skull, brain) at each position in the grid. These values were derived using mappings of CT intensity, based on Hounsfield unit values, to corresponding values for medium density, speed of sound, and absorption using the porosity method, as performed by Legon et al.^10^

These rectilinear grids were then used to simulate LIFUP from a fixed simulated transducer on the subject’s head. We modeled a 72mm diameter single-element transducer positioned in the grid based on the transformed transducer point. We defined a focal point tracing forward 55 mm (the length of our transducer’s focal depth from its membrane) along the line from the transducer point to the pallidal target point. Wave propagation from the transducer converging toward this focal point was simulated in the grid. We modeled the pressure wave emitted from the transducer as a three-dimensional sinusoidal wave using parameters obtained from water tank experiments (Acetera), specifically including a pressure at the face of the transducer of 1.0558 MPa and a carrier frequency of 650kHz. We simulated the propagation of this pressure wave from the transducer by solving coupled 1st-order nonlinear differential equations on the rectilinear grids of values representing density and speed of sound. The combination of these equations yields the generalized form of the Westervelt equation^61^. We solved this system of equations using k-Wave^26^. The maximal pressure and particle velocity values encountered at every voxel over time in the rectilinear grid simulation domain were recorded for analysis.

This scheme was simulated for a length of time corresponding to one pulse of the mode 2 parameter set used here (see ultrasound waveform for a complete description of parameter sets). In the mode 2 (10Hz Pulse Repetition Frequency) scheme, our pulse is on for 0.005 seconds and off for 0.095 seconds (0.1 seconds Pulse Period for a duty cycle of 5%). We thus simulated a duration of 0.005 seconds, or 3250 cycles at 650kHz (Carrier Frequency). We performed the simulation twice, first using the CT-derived head model and then using a free-water model, applying the same trajectory in both cases. For each simulation, the peak intensity inside the brain (the majority of energy deposition is in the skull when it is present) as well as its voxel coordinate were captured. We calculated the distance between the simulated point of highest intensity in free water and through bone in order to capture the degree of refraction expected to occur due to propagation through skull. Similarly, the peak intensity in both free-water and CT simulations were compared in order to quantify the degree of peak pressure attenuation expected due to skull. The cumulative energy deposition over time was calculated for the voxel of peak intensity for the through-skull simulation. Energy deposition over time was highly linear, suggesting that the shorter pulse width used here, 0.5ms, in terms of energy deposition, behaves the same as the longer pulse (see Figure S9) and that the results presented generalize to both pulse widths.

### BOLD Data Analysis

fMRI as well as ASL preprocessing and analysis was conducted using FSL (FMRIB Software Library v6.0.1)^62^ with in-house Bash shell scripts. In addition, second level data analysis was performed using JASP. JASP Team (2019). JASP (Version 0.11.1).

#### Preprocessing

Prior to analysis, preprocessing was performed including brain extraction (optibet^63^), spatial smoothing (using a Gaussian kernel of 5 mm full-width half-max), slice timing correction (Fourier-space time-series phase-shifting), highpass temporal filtering (Gaussian-weighted) at .01 Hz, and motion correction (MCFLIRT)^62,64^. None of the collected BOLD data exhibited motion greater than 3 mm translation or 3 degrees of rotation. Registration from functional to structural space for each subject was performed using FSL’s Boundary-Based Registration (BBR)^65^, with the exception of two subjects for which this failed; functional to structural space registrations for these subjects were performed using FSL FLIRT^62,64^ (Normal Search 12 DOF). Registration of functional to standard space for each run was performed using FSL FLIRT^62,64^ (Normal Search 12 DOF) and FSL’s nonlinear registration tool, FSL FNIRT^62^ (Warp resolution of 10 mm). Registration from structural to functional space as well as functional to standard space was confirmed visually for each run.

### BOLD – Whole Brain

LIFU-BOLD data sequences were first analyzed employing a univariate general linear model (GLM) approach^66^ including a pre-whitening correction for autocorrelation (FILM). For each LIFU-BOLD sequence for each participant, a univariate analysis was conducted using a single “task” regressor—onset time of 30s blocks of LIFU administration; moreover, 24 extended motion regressors were employed, including motion in 6 directions as well as first derivatives, second derivatives, and their difference. Thus, here, “baseline” refers to inter-sonication periods where no LIFU is applied. For each BOLD sequence, we computed 2 contrasts: LIFU > no LIFU and LIFU < no LIFU. Data from LIFU Mode 1 and data from LIFU Mode 2 were aggregated respectively and assessed statistically using a mixed effects FLAME 1 + 2 model. At level 2, these results were aggregated between runs of the same LIFU parameter set while the following contrasts were calculated on a per-subject basis: Mean LIFU Mode 1, Mean LIFU Mode 2, LIFU Mode 1 – LIFU Mode 2, LIFU Mode 2 – LIFU Mode 1, LIFU Mode 1 + LIFU Mode 2. At level 3, data were aggregated between subjects. Data were cluster corrected for multiple comparisons using a cluster-level threshold of z > 3.09 (corrected p < .05). A separate level 3 analysis was conducted with cluster correction at z > 2.57 (corrected p < .05).

### BOLD – ROI

Based on the trajectory of our ultrasound beam, three subcortical ROI’s (Left Putamen, Left Globus Pallidus and the Left Thalamus) were selected to determine if FUS modulated the BOLD signal in targeted regions and/or in adjacent regions. For each subject, masks for each region were created by using FSL’s automatic segmentation for subcortical nuclei (FIRST)^62^ for subcortical extraction on high resolution T1 images. Each subcortical ROI was binarized and confirmed for correct extraction visually. For each statistical z-score map produced in the GLM contrast of no LIFU blocks (see Block Design above) with LIFU blocks (LIFU > no LIFU), voxel-wise z-scores were averaged inside the area of each ROI using FSL’s tool for extracting the mean selected voxels in a 3D image (fslmeants)^62^. Registration via FSL’s linear registration tool (FLIRT^62,64^, Normal search, 12 DOF) of each z-score map to structural masks was confirmed visually. This leaves us with one number denoting the z-scored difference in BOLD between LIFU-on and baseline (LIFU-off) within each ROI for each LIFU run for each subject.

A 2 x 2 x 3 repeated measures ANOVA was performed with Parameter Set (Mode 1 or Mode 2), Run (Sonication 1 or Sonication 2 within session), and ROI (Left Putamen, Left GP, Left Thalamus) as factors. Follow-up 2 x 2 repeated measures ANOVAs were performed for each ROI with Parameter Set (Mode 1 or Mode 2), Run (Sonication 1 or Sonication 2 within session) as factors. In each of these follow-up ANOVAs, marginal means were computed for each parameter set and statistically assessed against zero in order to assess if an influence from each parameter set existed irrespective of the other. Šidák correction was used to correct for multiple comparisons across marginal mean test. Mauchly’s test of sphericity was found to not have been violated. The Shapiro Wilk test confirmed normality in the data. Several outliers existed in this data. A separate analysis was conducted with outliers excluded; this did not change results and so outliers were left in the data presented here.

In addition to frequentist testing, a 2 x 2 x 3 Bayesian repeated measures ANOVA was performed with Parameter Set, Run (Sonication 1 or Sonication 2 within session), and ROI (Left Putamen, Left GP, Left Thalamus) as factors. For all Bayesian t-tests employed throughout this study, a Cauchy distribution with a width parameter of 0.707 was used while, for all Bayesian ANOVAs, a r scale prior width of 0.5 was used; see JASP documentation for a discussion of Markov chain Monte Carlo (MCMC) settings, which are determined in a black-box manner by the JASP program. Follow-up 2 x 2 Bayesian repeated measures ANOVAs were also performed for each ROI. Bayes factors (BF) reported reflect the ratio of evidence for each alternative hypothesis (H1) against the null hypothesis(H0). A BF_10_ indicates the Bayes factor in favor of H1 over H0. A BF_10_ > 3 is widely considered as positive and substantial evidence for the alternative hypothesis^67^. BFIncluded denotes the evidence in support of the inclusion of effects (e.g., interaction terms) in the model over and above that of other terms. In order to estimate BF for marginal means, a separate analysis was performed in which runs were averaged. Bayesian one-sample one-sided t tests were conducted for each parameter for each ROI against zero in which the alternative hypothesis stated the data was below zero. Statistical analysis was performed in JASP.

### Connectivity Psychophysiological Interaction (PPI)

Psychophysiological analysis was conducted to determine which, if any, regions of the whole brain changed their connectivity with the left globus pallidus during 30s blocks of sonication as compared to non-sonication blocks. The same masks used for the BOLD ROI analyses (see Block Design BOLD ROI) were used here. This was done for data from each BOLD sequence (4 per subject) using FSL’s tool for extracting the time points of selected voxels in a 4D image (fslmeants)^62^. Using FSL FEAT, each LIFU-BOLD data sequence was first analyzed employing a multivariate general linear model (GLM) approach including our block design, the time series of the ROI, and importantly, their interaction. Finally, lower-level results from each LIFU mode were aggregated respectively using a mixed effects FLAME 1 + 2 model. Aggregated data were regressed on subject of origin. Data were cluster-corrected for multiple comparisons using a cluster-level threshold of z > 3.09 (corrected p < 0.05), as well as, in a separate third level analysis, z > 2.57 (corrected p < 0.05).

### Arterial Spin Labeling (ASL)

#### ASL – Whole Brain

In order to quantify the degree of blood perfusion throughout the brain at three time points for each subject visit— pre LIFU 1, post LIFU 1, and post LIFU 2—each ASL sequence was analyzed using standard processing methods in FSL Bayesian Inference for Arterial Spin Labeling MRI^68^ (BASIL), using the command-line tool oxford_asl. Analysis was set to conform to the assumptions concerning the kinetic model and T1 values outlined in the BASIL white paper^68^. The standard T1 value of atrial blood (T1b) was used (1.65). The inversion efficiency of pASL was set at 0.98. Estimation of bolus duration was disabled and supplied at 700ms. Spatial regularization^69^ as well as motion correction^62,64^ was used while artifact correction for ASL signal within the microvasculature was disabled . The resultant perfusion images were registered to standard space; the quality of these registrations was confirmed visually. At level 2, two linear and two “L” models were applied to the three estimates of perfusion derived from level one, for each mode (10Hz vs 100Hz) and for each subject. One linear model described a descending trend in blood perfusion through the session (1 0 −1) while the other, an ascending trend (−1 0 1). One “L” model described a descending trend in blood perfusion after LIFU 1 but decreasing no further (1 −.5 −.5) while the other, an ascending trend after LIFU 1 but then increasing no further (−1 .5 .5). At level 3, data was aggregated within respective modes and across subjects as well as statistically assessed using FSL randomise’s^70^ nonparametric permutation t test for voxel-based thresholding. A single-sample two-sided t test was run with threshold free cluster enhancement (TFCE)^33^, correcting the resultant *p*-values for multiple comparisons across space.

#### ASL – ROI

The ROI’s investigated in BOLD were also analyzed using ASL data. These included the Left Putamen, Left Globus Pallidus, and Left Thalamus. The same masks used for the BOLD ROI analyses (see Block Design BOLD ROI) were used here. Each perfusion image was registered to subject structural space using oxford_asl. Intensity of perfusion images were averaged inside the area of each ROI using FSL’s tool for extracting the mean selected voxels in a 3D image (fslmeants)^62^ for each perfusion image for each subject (16 images; 3 time points per 2 parameter sets).

The resultant intensity values were run in a 2 by 3 by 3 repeated measures ANOVA was run with Parameter Set (Mode 1 or Mode 2), Time Point (Pre LIFU 1, Post LIFU 1, Post LIFU 2) and ROI (Left Putamen, Left GP, Left Thalamus) as factors. A “Repeated” contrast was run for Time Point, which directly compares Pre LIFU 1 with Post LIFU 1 and Post LIFU 1 with Post LIFU 2. Follow-up 2 x 3, two-way repeated measures ANOVA’s were run for each ROI with Parameter Set (Mode 1 or Mode 2) and Time Point (Pre LIFU 1, Post LIFU 1, Post LIFU 2) as factors. A “Repeated” contrast was run for Time Point, which directly compares Pre LIFU 1 with Post LIFU 1 and Post LIFU 1 with Post LIFU 2 for each of these follow-up tests. Mauchly’s test of sphericity was found to be violated in several of these ANOVAs, thus the Greenhouse-Geisser correction for sphericity was applied to these. The Shapiro Wilk test confirmed normality in the data. No outliers were found in these data.

In addition to frequentist testing, a 2 by 3 by 3 Bayesian repeated measures ANOVA was run with Parameter Set (Mode 1 or Mode 2), Time Point (Pre LIFU 1, Post LIFU 1, Post LIFU 2) and ROI (Left Putamen, Left GP, Left Thalamus) as factors. In order to estimate BF for “Repeated” models, a follow-up comparison between Time Points was conducted. Statistical analysis was performed in JASP.

## Acknowledgements

This work was supported in part the Tiny Blue Dot foundation and by NIH grant R01-NS074980.

## Author Contributions

Joshua Cain authored the first draft of this manuscript, conducted data collection, and conducted neuroimaging analysis.

Martin Monti conceived of the experimental design, assisted with data collection, data analysis, and the writing of the first draft of the manuscript.

Shakthi Visagan and David Shattuck conducted acoustic simulations and assisted in the production of the figures relevant to the results of those simulations.

Micah Johnson Assisted with Bayesian statistical analysis.

Julia Crone provided guidance on neuroimaging analysis and on the theoretical implications of findings.

Robin Blades and Norman Spivak assisted in data collection.

All authors reviewed the manuscript and provided feedback before its submission.

## Data Availability Statement

Anonymized data are available from the first or last author and can be obtained through an MTA between the requesting PI/institution and the UCLA Technology Development Group (TDG).

## References

1. Bestmann, S. & Walsh, V. Transcranial electrical stimulation. Curr Biol 27, R1258–R1262 (2017).

2. Deng, Z.-D., Lisanby, S. H. & Peterchev, A. V. Electric field depth-focality tradeoff in transcranial magnetic stimulation: simulation comparison of 50 coil designs. Brain Stimul 6, 1–13 (2013).

3. Min, B.-K. et al. Focused ultrasound-mediated suppression of chemically-induced acute epileptic EEG activity. BMC Neuroscience 12, 23 (2011).

4. Yang, P. S. et al. Transcranial Focused Ultrasound to the Thalamus Is Associated with Reduced Extracellular GABA Levels in Rats. Neuropsychobiology 65, 153–160 (2012).

5. Yoo, S.-S., Kim, H., Min, B.-K. & Eric Franck, S. P. Transcranial Focused Ultrasound to the Thalamus Alters Anesthesia Time in Rats. Neuroreport 22, 783–787 (2011).

6. Min, B.-K. et al. Focused ultrasound modulates the level of cortical neurotransmitters: Potential as a new functional brain mapping technique. International Journal of Imaging Systems and Technology 21, 232–240 (2011).

7. Yoo, S.-S. et al. Focused ultrasound modulates region-specific brain activity. Neuroimage 56, 1267–1275 (2011).

8. Dallapiazza, R. F. et al. Noninvasive neuromodulation and thalamic mapping with low-intensity focused ultrasound. Journal of Neurosurgery 128, 875–884 (2017).

9. Folloni, D. et al. Manipulation of Subcortical and Deep Cortical Activity in the Primate Brain Using Transcranial Focused Ultrasound Stimulation. Neuron 101, 1109–1116.e5 (2019).

10. Legon, W., Ai, L., Bansal, P. & Mueller, J. K. Neuromodulation with single-element transcranial focused ultrasound in human thalamus. Hum Brain Mapp 39, 1995–2006 (2018).

11. Bystritsky, A. et al. A review of low-intensity focused ultrasound pulsation. Brain Stimulation 4, 125–136 (2011).

12. Kubanek, J. Neuromodulation with transcranial focused ultrasound. Neurosurg Focus 44, E14 (2018).

13. Lanciego, J. L., Luquin, N. & Obeso, J. A. Functional Neuroanatomy of the Basal Ganglia. Cold Spring Harb Perspect Med 2, (2012).

14. Schiff, N. D. Recovery of consciousness after brain injury: a mesocircuit hypothesis. Trends Neurosci 33, 1–9 (2010).

15. Qiu, M.-H., Yao, Q.-L., Vetrivelan, R., Chen, M. C. & Lu, J. Nigrostriatal Dopamine Acting on Globus Pallidus Regulates Sleep. Cereb Cortex 26, 1430–1439 (2016).

16. Yuan, X.-S. et al. Striatal adenosine A2A receptor neurons control active-period sleep via parvalbumin neurons in external globus pallidus. eLife 6, e29055 (2017).

17. Chen, M. C. et al. Identification of a direct GABAergic pallidocortical pathway in rodents. Eur J Neurosci 41, 748–759 (2015).

18. Saunders, A. et al. A direct GABAergic output from the basal ganglia to frontal cortex. Nature 521, 85–89 (2015).

19. Zheng, Z. S. & Monti, M. M. Thalamic and extra-thalamic connections of the Globus Pallidus in the human brain: The ultradirect pathway. bioRxiv 688283 (2019) doi:10.1101/688283.

20. Monti, M. M., Schnakers, C., Korb, A. S., Bystritsky, A. & Vespa, P. M. Non-Invasive Ultrasonic Thalamic Stimulation in Disorders of Consciousness after Severe Brain Injury: A First-in-Man Report. Brain Stimulation 9, 940–941 (2016).

21. Greenberg, B. D. et al. Deep brain stimulation of the ventral internal capsule/ventral striatum for obsessive-compulsive disorder: worldwide experience. Molecular Psychiatry 15, 64–79 (2010).

22. Schrock, L. E. et al. Tourette syndrome deep brain stimulation: A review and updated recommendations. Movement Disorders 30, 448–471 (2015).

23. Brunoni, A. R. et al. Transcranial direct current stimulation for acute major depressive episodes: meta-analysis of individual patient data. Br J Psychiatry 208, 522–531 (2016).

24. Agnesi, F., Johnson, M. D. & Vitek, J. L. Chapter 4 - Deep brain stimulation: how does it work? in Handbook of Clinical Neurology (eds. Lozano, A. M. & Hallett, M.) vol. 116 39–54 (Elsevier, 2013).

25. Edwards, T. C., Zrinzo, L., Limousin, P. & Foltynie, T. Deep brain stimulation in the treatment of chorea. Movement Disorders 27, 357–363 (2012).

26. Treeby, B. E. & Cox, B. T. k-Wave: MATLAB toolbox for the simulation and reconstruction of photoacoustic wave fields. J Biomed Opt 15, 021314 (2010).

27. Duck, F. A. Medical and non-medical protection standards for ultrasound and infrasound. Progress in Biophysics and Molecular Biology 93, 176–191 (2007).

28. Mueller, J. K., Ai, L., Bansal, P. & Legon, W. Numerical evaluation of the skull for human neuromodulation with transcranial focused ultrasound. J Neural Eng 14, 066012 (2017).

29. Spitzer, V. M. & Whitlock, D. G. The visible human dataset: The anatomical platform for human simulation. The Anatomical Record 253, 49–57 (1998).

30. Brinker, S. T., Preiswerk, F., McDannold, N. J., Parker, K. L. & Mariano, T. Y. Virtual Brain Projection for Evaluating Trans-skull Beam Behavior of Transcranial Ultrasound Devices. Ultrasound in Medicine & Biology 45, 1850–1856 (2019).

31. Eklund, A., Nichols, T. E. & Knutsson, H. Cluster failure: Why fMRI inferences for spatial extent have inflated false-positive rates. Proc. Natl. Acad. Sci. U.S.A. 113, 7900–7905 (2016).

32. Woo, C.-W., Krishnan, A. & Wager, T. D. Cluster-extent based thresholding in fMRI analyses: pitfalls and recommendations. Neuroimage 91, 412–419 (2014).

33. Smith, S. M. & Nichols, T. E. Threshold-free cluster enhancement: addressing problems of smoothing, threshold dependence and localisation in cluster inference. Neuroimage 44, 83–98 (2009).

34. Smith, S. M. et al. Advances in functional and structural MR image analysis and implementation as FSL. Neuroimage 23 Suppl 1, S208–219 (2004).

35. Ai, L., Mueller, J. K., Grant, A., Eryaman, Y. & Legon, W. Transcranial focused ultrasound for BOLD fMRI signal modulation in humans. in 2016 38th Annual International Conference of the IEEE Engineering in Medicine and Biology Society (EMBC) 1758–1761 (IEEE, 2016). doi:10.1109/EMBC.2016.7591057.

36. Pichardo, S., Milleret, R., Curiel, L., Pichot, O. & Chapelon, J.-Y. In vitro experimental study on the treatment of superficial venous insufficiency with high-intensity focused ultrasound. Ultrasound Med Biol 32, 883–891 (2006).

37. Sato, T., Shapiro, M. G. & Tsao, D. Y. Ultrasonic Neuromodulation Causes Widespread Cortical Activation via an Indirect Auditory Mechanism. Neuron 98, 1031–1041.e5 (2018).

38. Crone, J. S., Lutkenhoff, E. S., Bio, B. J., Laureys, S. & Monti, M. M. Testing Proposed Neuronal Models of Effective Connectivity Within the Cortico-basal Ganglia-thalamo-cortical Loop During Loss of Consciousness. Cereb Cortex 27, 2727–2738 (2017).

39. Bestmann, S. & Feredoes, E. Combined neurostimulation and neuroimaging in cognitive neuroscience: past, present, and future. Ann. N. Y. Acad. Sci. 1296, 11–30 (2013).

40. Verhagen, L. et al. Offline impact of transcranial focused ultrasound on cortical activation in primates. eLife 8, e40541 (2019).

41. Fouragnan, E. F. et al. The macaque anterior cingulate cortex translates counterfactual choice value into actual behavioral change. Nat Neurosci 22, 797–808 (2019).

42. Min, H.-K. et al. Deep brain stimulation induces BOLD activation in motor and non-motor networks: An fMRI comparison study of STN and EN/GPi DBS in large animals. NeuroImage 63, 1408–1420 (2012).

43. Jech, R. Functional Imaging of Deep Brain Stimulation: fMRI, SPECT, and PET. in Deep Brain Stimulation in Neurological and Psychiatric Disorders (eds. Tarsy, D., Vitek, J. L., Starr, P. A. & Okun, M. S.) 179–201 (Humana Press, 2008). doi:10.1007/978-1-59745-360-8_9.

44. Aguirre, G. K., Detre, J. A., Zarahn, E. & Alsop, D. C. Experimental Design and the Relative Sensitivity of BOLD and Perfusion fMRI. NeuroImage 15, 488–500 (2002).

45. King, R. L., Brown, J. R., Newsome, W. T. & Pauly, K. B. Effective Parameters for Ultrasound-Induced In Vivo Neurostimulation. Ultrasound in Medicine & Biology 39, 312–331 (2013).

46. Bystritsky, A. & Korb, A. S. A Review of Low-Intensity Transcranial Focused Ultrasound for Clinical Applications. Curr Behav Neurosci Rep 2, 60–66 (2015).

47. Smith, J. C., Paulson, E. S., Cook, D. B., Verber, M. D. & Tian, Q. Detecting changes in human cerebral blood flow after acute exercise using arterial spin labeling: Implications for fMRI. Journal of Neuroscience Methods 191, 258–262 (2010).

48. Chang, C. & Glover, G. H. Relationship between respiration, end-tidal CO2, and BOLD signals in resting-state fMRI. Neuroimage 47, 1381–1393 (2009).

49. Hajjar Ihab, Zhao Peng, Alsop David & Novak Vera. Hypertension and Cerebral Vasoreactivity. Hypertension 56, 859–864 (2010).

50. Plaksin, M., Kimmel, E. & Shoham, S. Cell-Type-Selective Effects of Intramembrane Cavitation as a Unifying Theoretical Framework for Ultrasonic Neuromodulation. eNeuro 3, (2016).

51. Tyler, W. J. Noninvasive Neuromodulation with Ultrasound? A Continuum Mechanics Hypothesis. Neuroscientist 17, 25–36 (2011).

52. Plaksin, M., Shoham, S. & Kimmel, E. Intramembrane Cavitation as a Predictive Bio-Piezoelectric Mechanism for Ultrasonic Brain Stimulation. Phys. Rev. X 4, 011004 (2014).

53. Lutkenhoff, E. S. et al. Thalamic and extrathalamic mechanisms of consciousness after severe brain injury. Annals of Neurology 78, 68–76 (2015).

54. Fenoy, A. J. & Simpson, R. K. Risks of common complications in deep brain stimulation surgery: management and avoidance. J. Neurosurg. 120, 132–139 (2014).

55. Schnakers, C. & Monti, M. M. Disorders of consciousness after severe brain injury: therapeutic options. Current Opinion in Neurology 30, 573–579 (2017).

56. Izadifar, Z., Izadifar, Z., Chapman, D. & Babyn, P. An Introduction to High Intensity Focused Ultrasound: Systematic Review on Principles, Devices, and Clinical Applications. J Clin Med 9, (2020).

57. Powers, W. J., Grubb, R. L., Darriet, D. & Raichle, M. E. Cerebral blood flow and cerebral metabolic rate of oxygen requirements for cerebral function and viability in humans. J. Cereb. Blood Flow Metab. 5, 600–608 (1985).

58. Legon, W. et al. Transcranial focused ultrasound modulates the activity of primary somatosensory cortex in humans. Nat. Neurosci. 17, 322–329 (2014).

59. Sanguinetti, J. L. et al. Transcranial Focused Ultrasound to the Right Prefrontal Cortex Improves Mood and Alters Functional Connectivity in Humans. Front. Hum. Neurosci. 14, (2020).

60. Lee, W. et al. Non-invasive transmission of sensorimotor information in humans using an EEG/focused ultrasound brain-to-brain interface. PLOS ONE 12, e0178476 (2017).

61. Wise, E. S. & Treeby, B. E. Full-wave nonlinear ultrasound simulation in an axisymmetric coordinate system using the discrete sine and cosine transforms. in 2013 IEEE International Ultrasonics Symposium (IUS) 1374–1377 (2013). doi:10.1109/ULTSYM.2013.0349.

62. Jenkinson, M., Beckmann, C. F., Behrens, T. E. J., Woolrich, M. W. & Smith, S. M. FSL. NeuroImage 62, 782–790 (2012).

63. Lutkenhoff, E. S. et al. Optimized Brain Extraction for Pathological Brains (optiBET). PLOS ONE 9, e115551 (2014).

64. Jenkinson, M., Bannister, P., Brady, M. & Smith, S. Improved optimization for the robust and accurate linear registration and motion correction of brain images. Neuroimage 17, 825–841 (2002).

65. Greve, D. N. & Fischl, B. Accurate and Robust Brain Image Alignment using Boundary-based Registration. Neuroimage 48, 63–72 (2009).

66. Monti, M. M. Statistical Analysis of fMRI Time-Series: A Critical Review of the GLM Approach. Front. Hum. Neurosci. 5, (2011).

67. Kass, R. E. & Raftery, A. E. Bayes Factors. Journal of the American Statistical Association 90, 773–795 (1995).

68. Chappell, M. A., Groves, A. R., Whitcher, B. & Woolrich, M. W. Variational Bayesian Inference for a Nonlinear Forward Model. IEEE TRANSACTIONS ON SIGNAL PROCESSING 57, (2009).

69. Groves, A. R., Chappell, M. A. & Woolrich, M. W. Combined spatial and non-spatial prior for inference on MRI time-series. Neuroimage 45, 795–809 (2009).

70. Winkler, A. M., Ridgway, G. R., Webster, M. A., Smith, S. M. & Nichols, T. E. Permutation inference for the general linear model. Neuroimage 92, 381–397 (2014).

