## Supplementary Information for "Real Time and Delayed Effects of Subcortical Low Intensity Focused Ultrasound"

### SUPPLEMENT

| Region | Voxels | Z-Max | Z-Max (MNI) | | |
|---|---|---|---|---|---|
| | | | x | y | z |
| LIFU Mode 1 – Baseline | | | | | |
| *Midline Precentral Ctx. | 4127 | 5.01 | 4 | -18 | 76 |
| *Posterior Cingulate Ctx. | 855 | 4.1 | 6 | -40 | 0 |
| *Heschel's Gyrus | 484 | 3.71 | 60 | -6 | 8 |
| *Frontal Polar Ctx. | 457 | 4.15 | -2 | 62 | 32 |
| Frontal Medial Ctx. | 234 | 3.6 | -4 | 46 | -16 |
| LIFU Mode 1 – LIFU Mode 2 | | | | | |
| Midline Postcentral Gyrus | 510 | 4.31 | 38 | -34 | 64 |
| Midline Precentral Gyrus | 245 | 4.04 | -4 | -30 | 74 |
| Posterior Cingulate Ctx. | 218 | 3.87 | 8 | -40 | 0 |

**Table S1: BOLD Changes in Whole-Brain During 100Hz Sonication Compared to Baseline, 10Hz Sonication.** Significant clusters as defined by a cluster significance level of $p < 0.05$ with a cluster defining threshold (CDT) of $p < 0.005$. Regions that also survived a more conservative CDT of $p < 0.001$ are marked with an asterisk *.

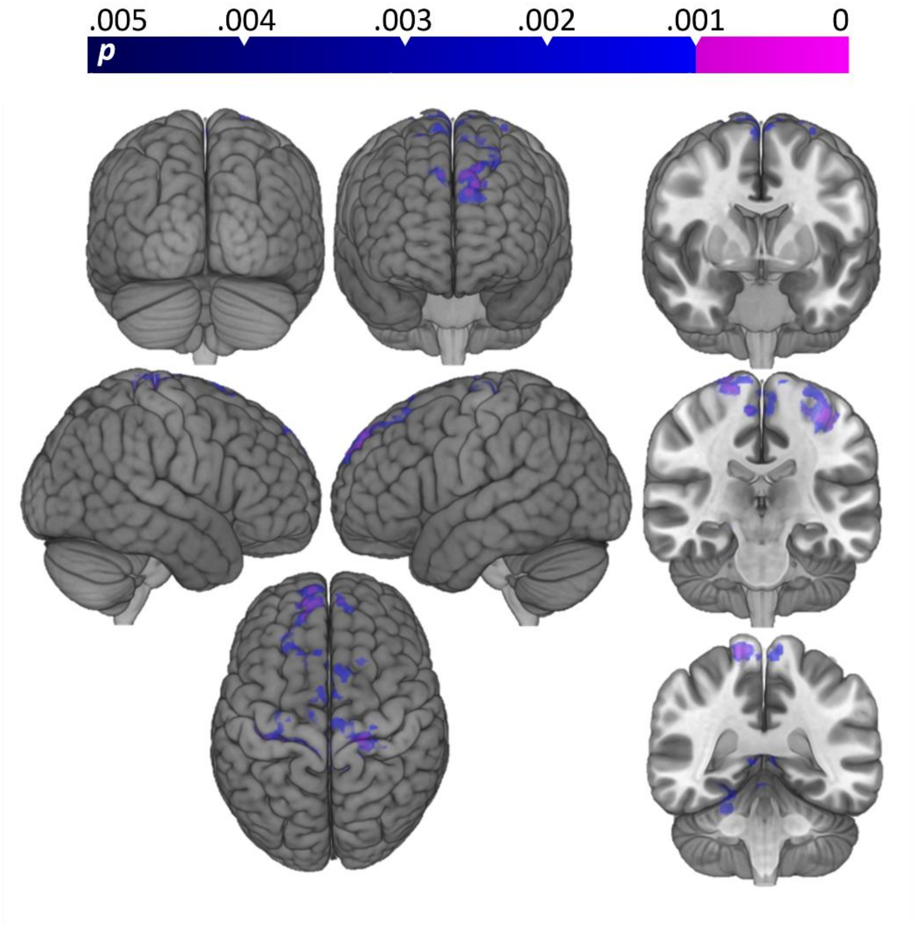


**Figure S1: Aggregating LIFU Mode 1 and LIFU Mode 2 data.** Shown here are the results of modeling our block design on data captured during both LIFU modes (Mixed Effects FLAME 1+2; cluster significance: $p < 0.05$; cluster defining threshold: $p < 0.005$ (blue), $p < 0.001$(violet)). Despite the boost in statistical power theoretically provided by doubling the data included, no new clusters are added to the results obtained when analyzing LIFU Mode 1 alone.

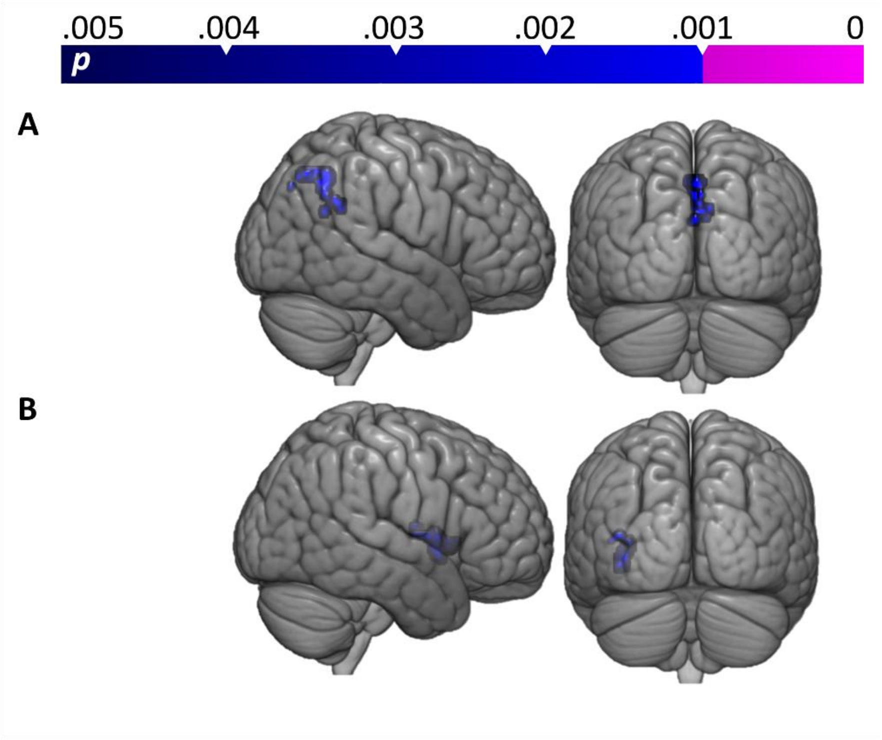


**Figure S2: Comparisons between Run1 and Run2 for LIFU in Mode 1 (PRF = 100Hz).** Shown here is a statistical map (Mixed Effects FLAME 1+2; cluster significance: $p < 0.05$; cluster defining threshold: $p < 0.005$ (blue), $p < 0.001$(violet)) of results obtained when subtracting successive LIFU runs (two data sets were collected per session) during Mode 1 (PRF = 100Hz) LIFU. **A)** Run1 -Run2 **B)** Run2-Run1. Significant voxels denote regions of relatively decreased BOLD (i.e., more suppression of BOLD signal from baseline). Minor differences exist, suggesting no major sensitization or habituation to LIFU's influence.

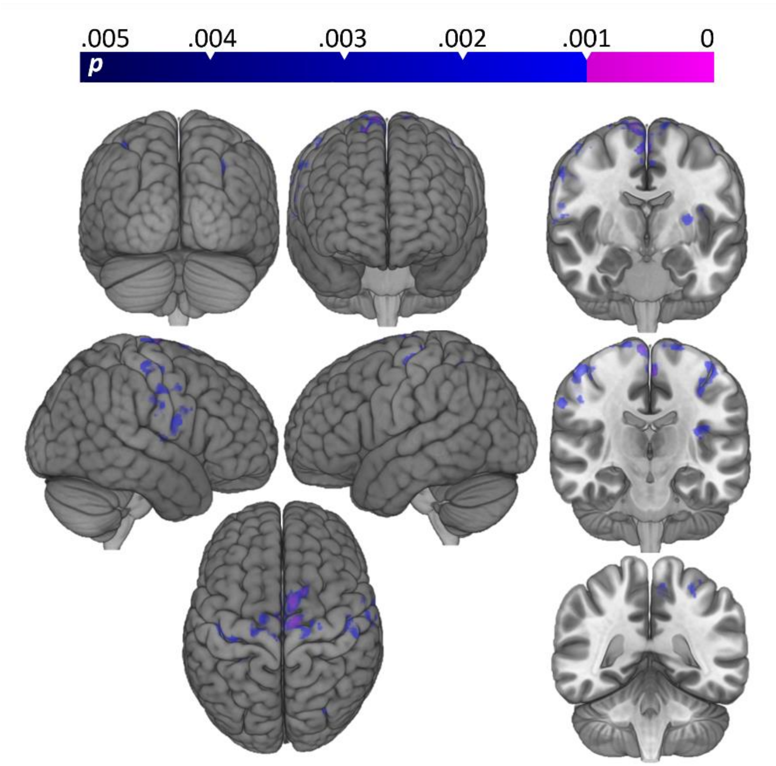


**Figure S3: Comparisons between Run1 and Run2 for LIFU in Mode 2 (PRF = 10Hz).** Shown here is a statistical map (Mixed Effects FLAME 1+2; cluster significance: $p < 0.05$; cluster defining threshold: $p < 0.005$ (blue), $p < 0.001$(violet)) of results obtained when subtracting successive LIFU runs (two data sets were collected per session) during Mode 2 (PRF = 10Hz) LIFU**.** Significant voxels denote regions of relatively decreased BOLD (i.e., more suppression of BOLD signal from baseline). We find a greater inhibition of BOLD signal in Run 2 compared to Run 1, in regions generally in line with inhibition found when comparing LIFU Mode 1 to baseline, suggesting that perhaps a sensitization occurs during LIFU in Mode 2 over time.

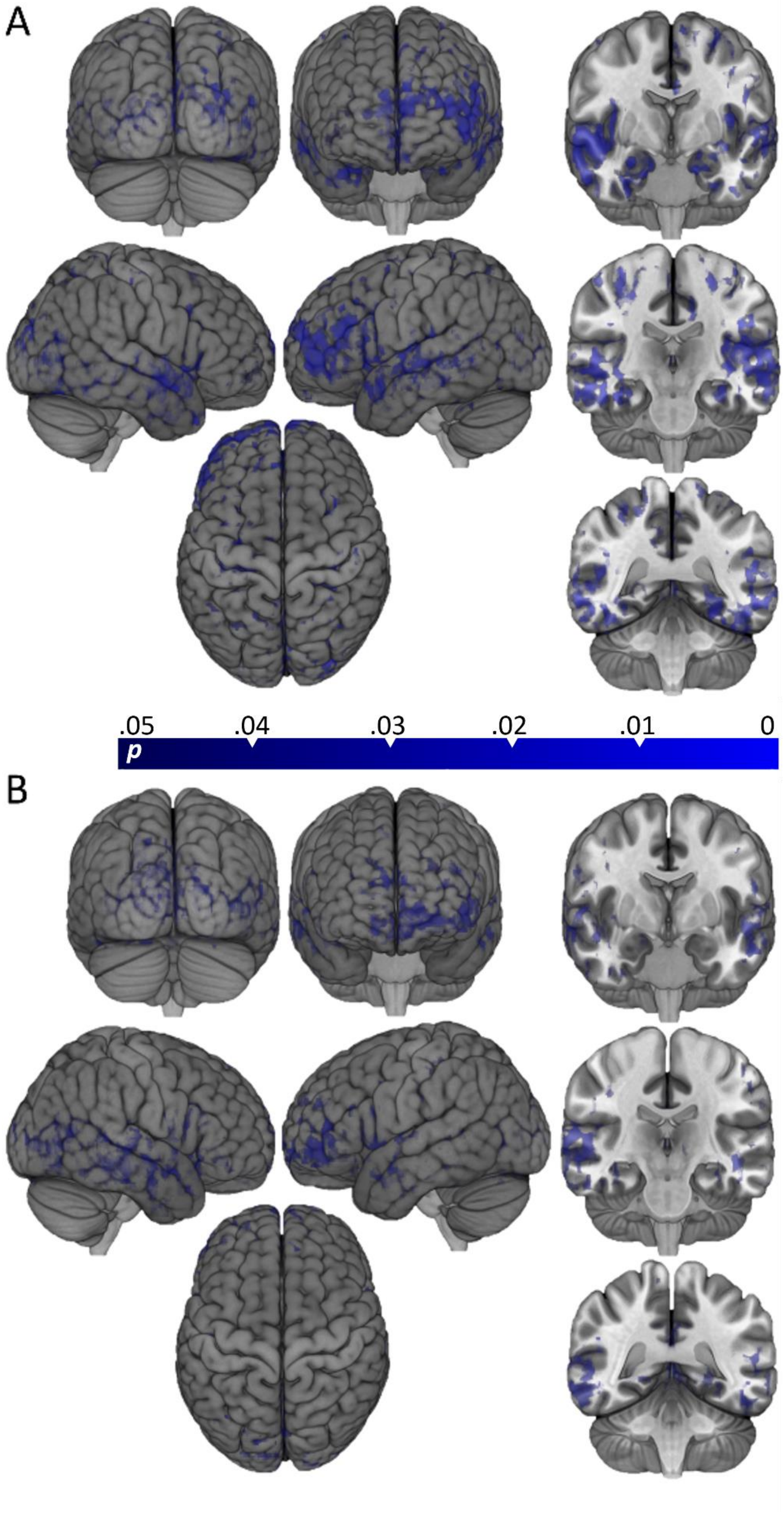
A
.05
.04
.03
.02
.01
0
p
B

**Figure S4: Whole-Brain Perfusion Linear Model.** Statistical maps of Blood Perfusion resulting from ASL analysis for all subjects (n=16), including 3 total ASL captures per subject per parameter set—taken immediately before and after each pallidal LIFU in **A)** Mode 1 (100Hz PRF; 5ms PW) and **B)** Mode 2 (10Hz PRF; 0.5ms PW). Color voxels indicate those that fit a linear shaped model such that activity decreased following sonication 1 and decreased further following sonication 2. The colored bar indicates the p-value window with $p < 0.05$. No increase in blood perfusion at these parameters is indicated because none was found. No subtraction between parameters is shown because none was found.

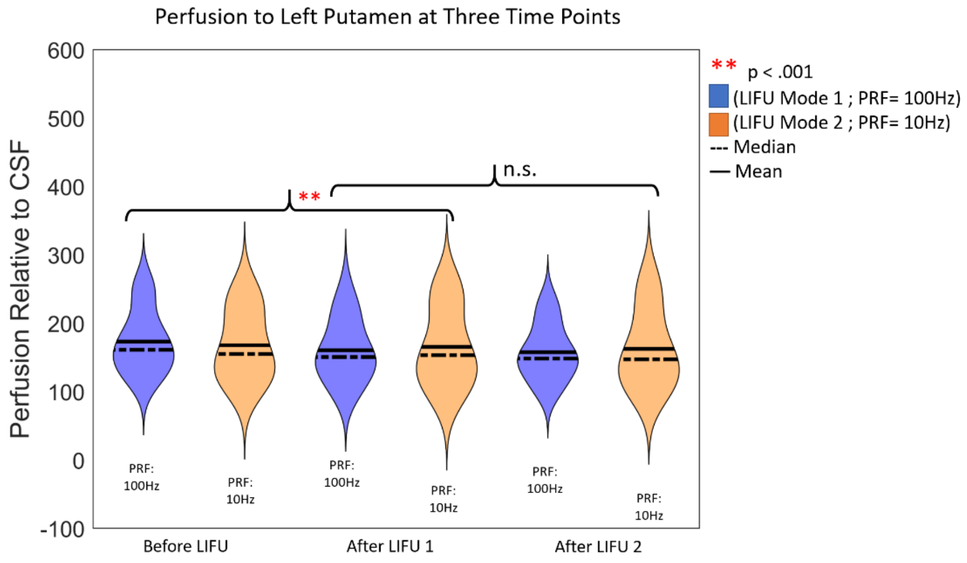


**Figure S5: ASL ROI Putamen.** The perfusion signal from the Left Putamen at three time points: Before LIFU 1, After LIFU 1, and After LIFU 2. A main effect of Time was found, while a significant decrease in perfusion was found following LIFU 1. However, perfusion was not found to decrease further following LIFU 2.

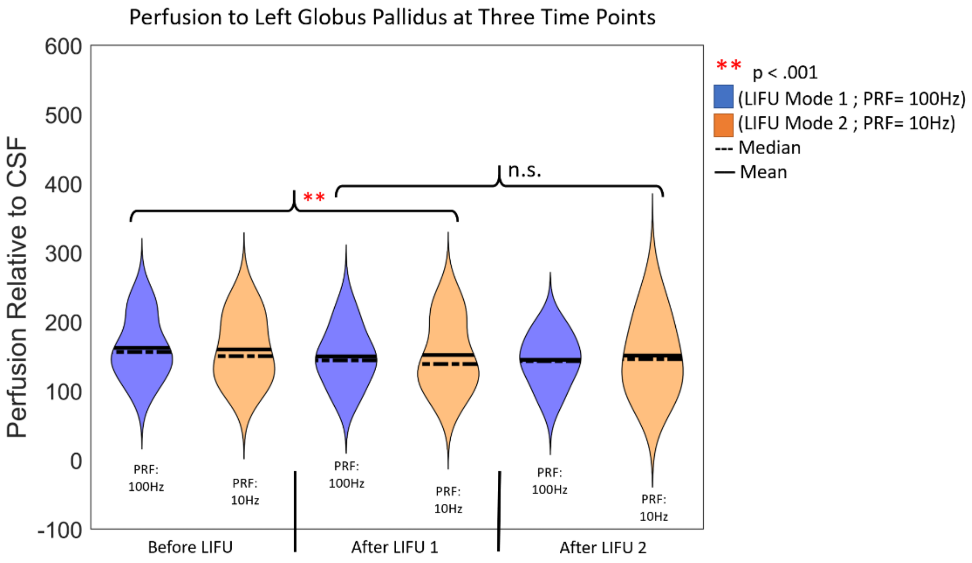


**Figure S6: ASL ROI Left Globus Pallidus.** The perfusion signal from the Left Globus Pallidus at three time points: Before LIFU 1, After LIFU 1, and After LIFU 2. A main effect of Time was found, while a significant decrease in perfusion was found following LIFU 1. However, perfusion was not found to decrease further following LIFU 2.

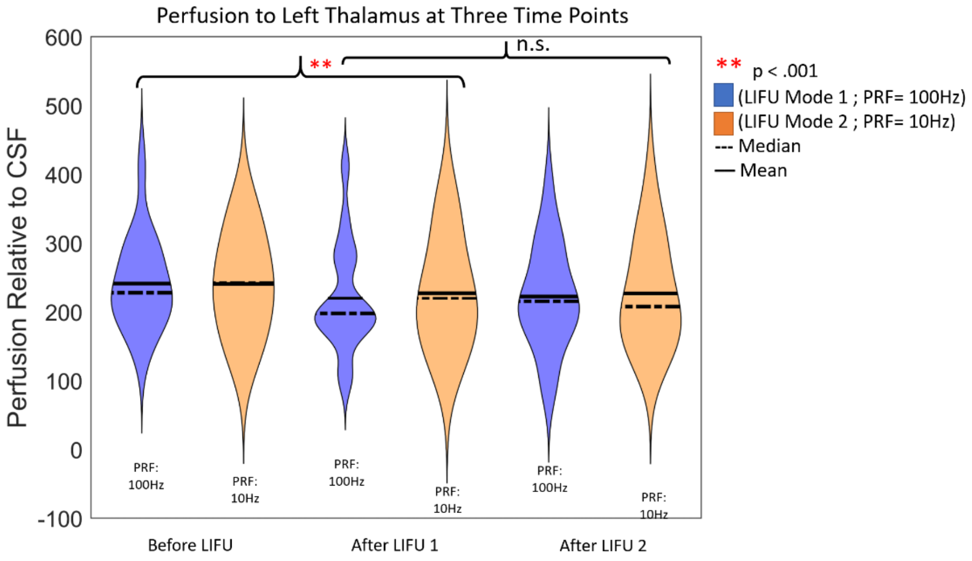

**Figure S7: ASL ROI Left Thalamus.** The perfusion signal from the Left Thalamus at three time points: Before LIFU 1, After LIFU 1, and After LIFU 2. A main effect of Time was found while a significant decrease in perfusion was found following LIFU 1. However, perfusion was not found tot decrease further following LIFU 2.

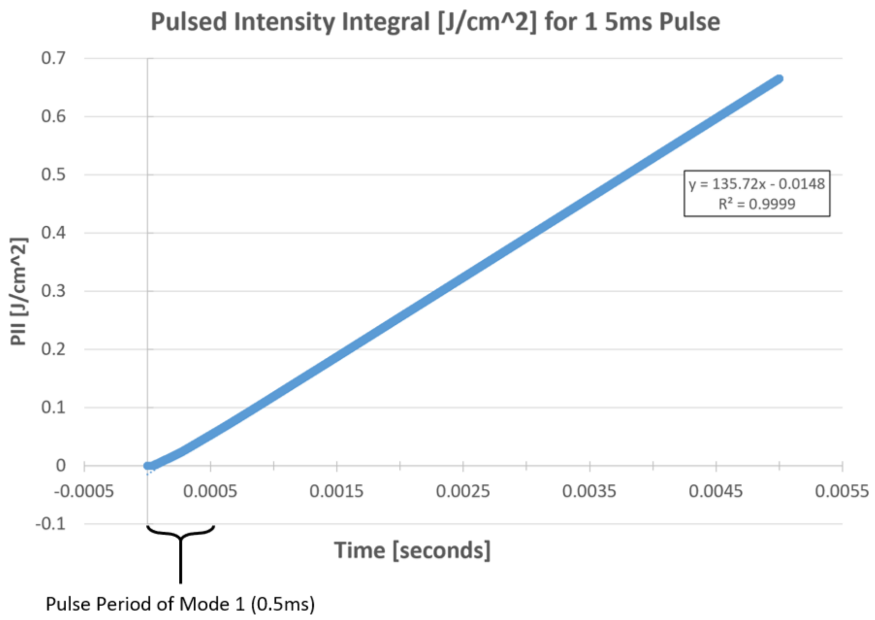


**Figure S8: Cumulative Pressure Over Time.** Results from monitoring the pressure at the voxel of maximum intensity (see Figure 2) when simulating a 5ms pulse over the full course of that pulse. The simulation scheme is the through-skull scheme described in Methods. As demonstrated by an $R^2$ of effectively 1, the pressure experienced by the brain appears highly linear over the time period of this pulse. This suggests a highly similar pressure distribution over-time for both pulse lengths used here. Note the length of the shorter pulse (0.5ms) represented here.
